# Hairy root transformation system as a tool for CRISPR/Cas9-directed genome editing in oilseed rape (*Brassica napus*)

**DOI:** 10.1101/2022.04.07.487540

**Authors:** Veronika Jedličková, Kateřina Mácová, Marie Štefková, Jan Butula, Jana Staveníková, Marek Sedláček, Hélène S. Robert

## Abstract

Our study examined the mutation efficiency of the CRISPR/Cas9 method for tryptophan aminotransferase *BnaTAA1* genes involved in the auxin biosynthesis pathway. We made nine CRISPR/Cas9 constructs with various promoters driving the expression of a Cas9 from *Staphylococcus aureus* (SaCas9) or a plant codon-optimized *Streptococcus pyogenes* Cas9 (pcoCas9). We developed a fast and efficient system for evaluating the variety and frequency of mutations caused by each construct using *Brassica napus* hairy roots. We showed that pcoCas9 is more efficient in mutating the targeted loci than SaCas9 and the presence of the NLS signal enhanced the chance of mutagenesis by 25%. The mutations were studied further in regenerated lines, and we determined the *BnaTAA1* gene expression and heritability of the gene modifications in transgenic plants. Hairy root transformation combined with CRISPR/Cas9-mediated gene editing represents a fast and straightforward system for studying target gene function in the important oilseed crop *Brassica napus*.

**One-sentence summary:** The hairy root transformation system of *Brassica napus* generates stable transformants and is a tool for efficiently identifying CRISPR/Cas9-induced genome editing.

## Introduction

The infection of *Agrobacterium* strains harboring a hairy root-inducing (*Ri*) plasmid causes an abnormal rooting on its plant hosts. After an agrobacterial infection at wounded sites, a T-DNA from the *Ri* plasmid is transferred to the host cells and stably integrated into the plant genome. Subsequently, the expression of T-DNA genes leads to the induction of hairy roots (Christey, 2001). The development of hairy roots is controlled mainly by root oncogenic loci (*rol*) genes (White et al., 1985, Cardarelli et al., 1987). *Ri* plasmids are classified according to the types of the T-DNA-encoded opine induced in hairy roots: cucumopine, mannopine, and agropine. For the latter, *Ri* plasmids carry two T-DNA regions, right and left T-DNA (TR-DNA and TL-DNA, respectively), which can independently integrate into the plant genome. For cucumopine and mannopine types, the T-DNA consists of a single region (Gelvin, 1990). The virulence (*vir*) region of the *Ri* plasmids contains numerous genes involved in the processing and transfer of the T-DNA from the bacteria to the plant cells (Gelvin, 2003). Agrobacterial strains carrying both *Ri* plasmid and artificial binary vector have been widely used for delivering foreign DNA into plant cells. Hairy root cultures have gained increasing attention as a system for the production of both recombinant proteins and valuable secondary metabolites with medicinal applications (Georgiev et al., 2012; Gutierrez-Valdes et al., 2020) or to investigate phytoremediation processes, i.e., using plants to clean up contaminated environments (Agostini et al., 2013).

Hairy root transformation has been recently discovered as an efficient tool for studying gene function using Clustered Regularly Interspaced Short Palindromic Repeats (CRISPR)/CRISPR-associated protein 9 (Cas9)-mediated gene editing in a wide range of plant taxa (reviewed in Kiryushkin et al., 2021). This bacterial system has been exploited to develop a powerful RNA-guided genome editing tool in various species. The mechanism resides in the employment of Cas9 endonuclease directed by a guide RNA (gRNA) to a specific sequence in the genome. A Cas9-gRNA complex binds and cuts the targeted DNA creating a double-strand break (DSB). Subsequently, the cell’s endogenous mechanisms repair the DSB, eventually introducing mutations into the target site (Deltcheva et al., 2011; Jinek et al., 2012; Doudna and Charpentier, 2014).

The CRISPR/Cas9 system has been employed to generate mutants in various crops, including rapeseed (*Brassica napus*). Since *B. napus* is allotetraploid species (2n = 38, AACC) formed by hybridization between *B. rapa* (2n = 20, AA) and *B. oleracea* (2n = 18, CC), it is necessary to mutate homoeologs from both subgenomes to alter a trait of interest. Targeted genetic mutations were introduced to improve rapeseed agronomic traits such as grain composition, plant architecture, or disease resistance (reviewed in Gocal et al., 2021). Most studies rely on traditional *A. tumefaciens*-based transformation of explants followed by regeneration of transgenic plants. An alternative protocol of rapeseed protoplast transfection with CRISPR/Cas9 vectors has been recently used, showing the protoplast regeneration as the main bottleneck of this approach (Lin et al., 2018; Li et al., 2021).

To avoid the lengthy explants transformation and regeneration process, we exploited hairy root cultures to study the gene-editing efficiency of various CRISPR/Cas9 vectors designed to target the auxin biosynthetic gene *TRYPTOPHAN AMINOTRANSFERASE* (*BnaTAA1*). Further, we regenerated T0 mutant plants from hairy root lines to study *BnaTAA1* gene expression and heritability of the gene modifications. Finally, we produced transgene-free and target gene-edited plants in the T1 generation.

## Results

### Selection of CRISPR/Cas9 constructs to target *BnaTAA1* genes for mutagenesis

We targeted *Brassica napus BnaTAA1* genes for mutagenesis to study auxin biosynthesis in oilseed rape. A BLAST search against the *Brassica napus* genome (B. napus v4.1, Genoscope) was performed using the *Arabidopsis thaliana TAA1* (*At1g70560*) coding sequence as a query to identify the *BnaTAA1* loci. We identified four genes (*BnaA02g14990D, BnaAnng22030D, BnaC02g19980D, BnaC06g43720D*) with 82 % to 86 % nucleotide sequence identity to *AtTAA1* and five genes (*BnaA09g31180D, BnaA09g31200D, BnaC05g18610D BnaA01g14030D, BnaC01g16530D*) representing *TAA-related 1* (*TAR1*) and *TAR2* (**Supplemental Figure S1**).

Proteins encoded by *BnaA02g14990D* (*BnaA02*.*TAA1*) and *BnaC02g19980D* (*BnaC02*.*TAA1*) share 98% amino acid identity, and both genes are expressed during seed development (PRJNA311067, GEO: GSE77637, Wan et al., 2017). *BnaAnng22030D* codes for a partial TAA1 protein with predicted premature stop codon in exon III. We confirmed the occurrence of this stop codon by amplifying and sequencing the corresponding genomic locus (Genbank, OM687491). On the other hand, a gene prediction tool (Keller et al., 2011) suggested the presence of an alternative spliced variant, enabling reading frame preservation. The predicted protein from the altered coding sequence shares 90 % amino acid similarity with BnaA02.TAA1 and BnaC02.TAA1 over the full length of the sequences. Phylogenetic analysis shows that BnaAnng22030D represents a paralog of the BnaTAA1 family (**Supplemental Figure S1**). *BnaAnng22030D* and *BnaC06g43720D* are expressed at very low levels in oilseed rape compared to *BnaA02*.*TAA1* and *BnaC02*.*TAA1* (*Brassica* Expression Database, Chao et al., 2020).

Because plant transformation, mutant selection, and plant regeneration are laborious and time-consuming protocols, especially in crops, we compared the mutagenesis effectiveness of pcoCas9 and SaCas9 proteins on the rapeseed genome. The pcoCas9 (plant-codon optimized Cas9), derived from *Streptococcus pyogenes* Cas9 (SpCas9), was optimized for plant genomes and contains the potato IV2 intron (Li et al., 2013). The *SaCas9* gene is about 1 kb shorter than *pcoCas9*. Its sequence has also been codon-optimized for *A. thaliana* (Steinert et al., 2015). Furthermore, SaCas9 may be more effective for gene editing in Arabidopsis than SpCa9 (Wolter et al., 2018). Both pcoCas9 and SaCas9 require specific gRNA sequence due to their secondary structure and, therefore, recognize different protospacer adjacent motifs (PAM), NNGRRT for SaCas9 and NGG for pcoCas9, which provide variability in targets, specificity (reduced off-target mutations of SaCas9), and efficiency (Kaya et al., 2016). Using CRISPR-P v2 online tool, we designed two sets of two guides (1) targeting identical sequences in *BnaA02*.*TAA1* and *BnaC02*.*TAA1* to mutagenize both loci with one set of guides and (2) enabling the creation of a large deletion between the two targeted loci (**Figure 1, Supplemental Table S1**). The target sequence of SaCas9 guide2 in paralogous genes, *BnaAnng22030D* and *BnaC06g43720D*, contains a single SNP localized 9 bp upstream of the PAM sequence, consistent with a high off-target score of 0.722 calculated by CRISPR-P v2.0 (**Table 1**).

**Figure 1.**
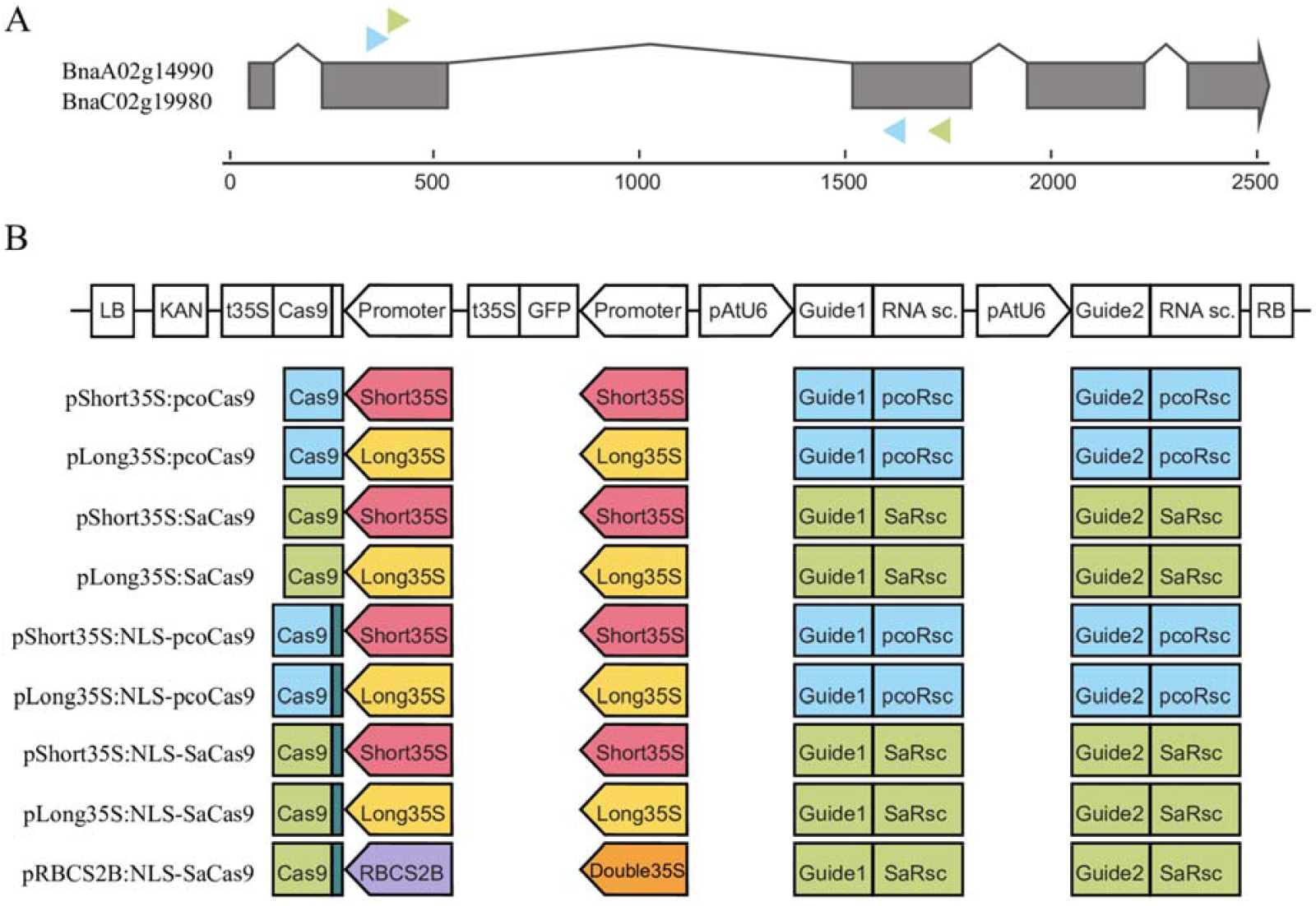
Design of the CRISPR/Cas9 constructs. A, Structure of the *BnaA02g14990D* (*BnaA02*.*TAA1*) and *BnaC02g19980D* (*BnaC02*.*TAA1*) genes with exons (grey boxes). The structure is identical for both genes. The colored arrowheads indicate the targeted position of the two gRNA for pcoCas9 (blue) and SaCas9 (green). B, Design of the nine CRISPR/Cas9 constructs. The T-DNA is composed in *cis* of a Kanamycin selective marker, the Cas9 cassette (promoter, Cas9 coding sequence, 35S terminator), the GFP cassette (promoter, GFP cds, 35S terminator), and the guide cassette (AtU6 promoter, gRNA, RNA scaffold). The types of promoters, Cas9, and guides are color-coded: pcoCas9 in blue, SaCas9 in green, NLS signal in dark green, short 35S promoter in red, long 35S promoter in yellow, double 35S promoter in orange, RBCS2B in purple, gRNA-RNA scaffold for pcoCas9 in blue, gRNA-RNA scaffold for SaCas9 in green.

**Table 1.**
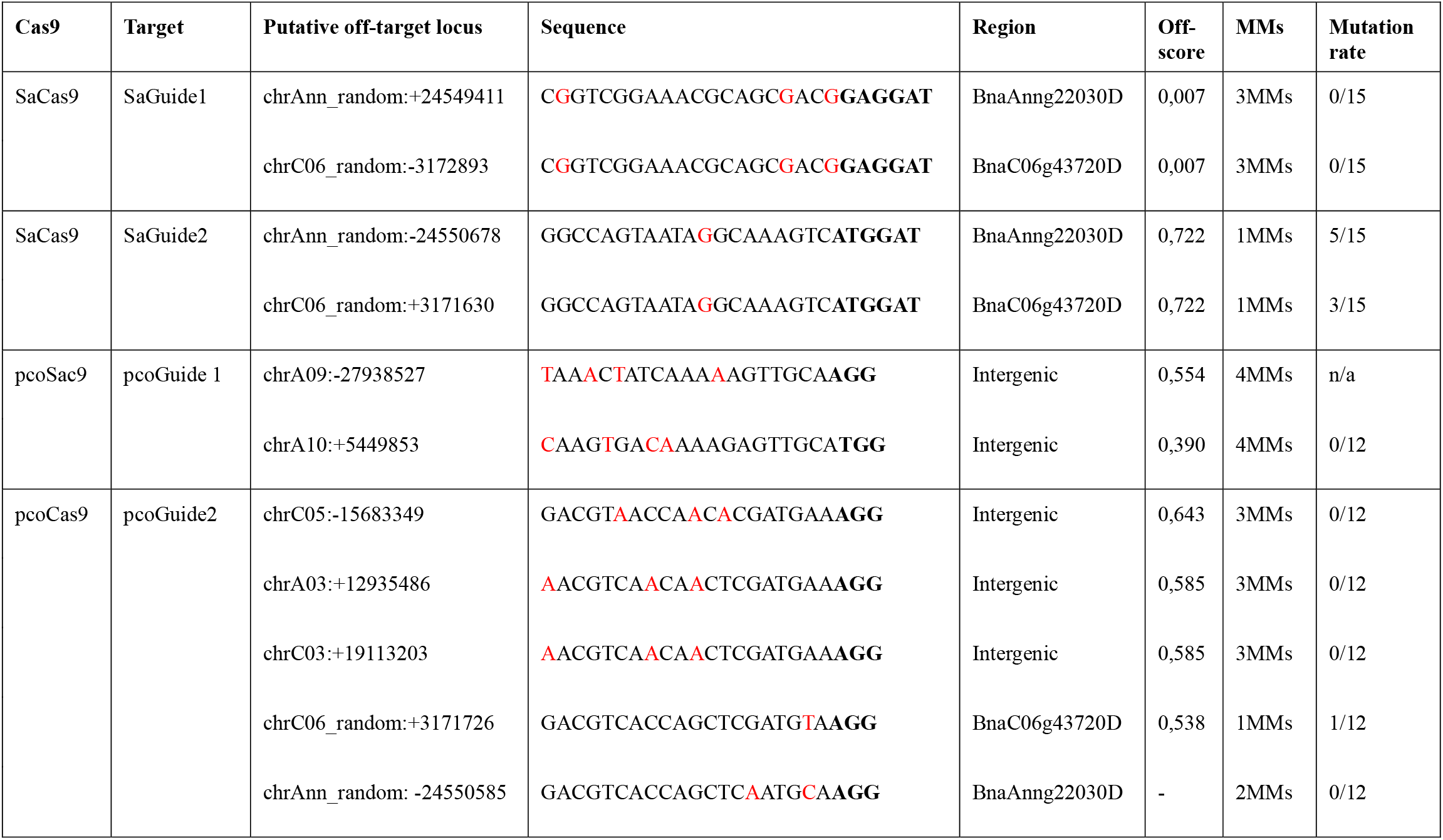
Detection of mutations at potential CRISPR/Cas9 off-target sites in the T0 plants. The off-score was predicted using the CRISPR-P v2.0 tool against the Darmor-bzh reference genome. The PAM motif is highlighted in bold, mismatched bases in red color. MMs, number of mismatches. Mutation rate, number of plants with mutation / number of tested plants.

The efficiency of the CRISPR/Cas9 system also depends on the promoter driving the *Cas9* expression. Plant-derived promoters can be as effective as the widely used Cauliflower Mosaic Virus *35S* promoter (Yan et al., 2015; Zhang et al., 2017). The choice of the promoter depends on the plant transformation methods (Yan et al., 2015). While a plant promoter driving *Cas9* expression in germline cells would be preferred in Arabidopsis, where floral dip transformation is the method of choice (Mao et al., 2016; Wang et al., 2015), ubiquitous promoters like *p35S* or *pUBIQUITIN* are favored in crop breeding where a strong *Cas9* expression in the vegetative tissues is necessary for transformation protocols using plant regeneration. Therefore, we compared two variations of the *35S* promoter (long and short). The *RIBULOSE BISPHOSPHATE CARBOXYLASE SMALL SUBU-NIT 2B* (*AtRBCS2B*) promoter was also selected to drive *SaCas9* expression. Finally, we tested whether the presence of a nuclear localization signal (SV40) would affect the efficiency of the whole system (Grützner et al., 2021).

We generated nine CRISPR/Cas9 constructs (**Figure 1, Supplemental Tables S2 and S3**), combining three promoters (long and short versions of *p35S* and *pRBCS2B*), with or without the SV40 nuclear targeting signal and with either pcoCas9 or SaCas9. The constructs are generated using a Modular Cloning system allowing the quick guides’ insertion into the constructs. In addition, each construct contains a green fluorescent (GFP) reporter to monitor the presence of the CRISPR/Cas9 transgene in plant tissues. These nine constructs were then transformed into *B. napus* using a hairy root transformation system.

### An efficient protocol for hairy root induction in *B. napus*

To develop a robust hairy root transformation system in *B. napus*, we tested three cultivars, namely DH12075, Topas DH4079, and Westar, for their transformation efficiency by an injection-based method. Hypocotyls of eighteen-day-old seedlings were injected with a suspension of agrobacterium containing the virulence *Ri* plasmid only. The first calli and hairy roots were detected two weeks after injection. The overall transformation efficiency (number of seedlings with emerging hairy roots over the number of injected seedlings) was calculated one month after the injection (**Supplemental Figure S2**). DH12075 has the highest transformation efficiency (97%), followed by Westar (84%) and Topas (42%), a cultivar recalcitrant to petiole-based transformation.

Because of its high transformation efficiency, the DH12075 cultivar was transformed with the CRISPR/Cas9 constructs to mutate the *BnaTAA1* genes. Roots emerging from the callus (1 – 3 per plant) were cut and transferred to a solid medium containing a selective agent (kanamycin). The GFP reporter in the CRISPR/Cas9 T-DNA eased the selection of transgenic roots by monitoring the presence of GFP fluorescence in the hairy roots. In total, 211 hairy root lines growing on kanamycin and expressing GFP were selected for further analysis.

### Evaluation of the mutagenesis efficiency of the CRISPR/Cas9 constructs in hairy roots

We analyzed 1 – 3 roots per independent transformant and 13 – 20 transformants per construct (16 – 30 transformed hairy roots per construct were analyzed) for the presence of a mutation in any of the four loci. All constructs were able to induce mutation with variable efficiency (**Figure 2, Supplemental Table S4**). The data shows that pcoCas9 (all promoters, NLS or not) is more efficient in mutating the targeted loci than SaCas9 (all promoters, NLS or not), with 83,05 % of mutated loci versus 47,98 %. We observed a slight efficiency increase with the long *35S* promoter (67,29 % mutated loci, compared to 58,7 % for short *35S* promoter). The presence of the NLS signal enhanced the chance of mutagenesis by 25 % (75,82 % with NLS-Cas9 versus 50,53 % with Cas9). The best construct based on the mutagenesis efficiency in this experiment is NLS-pcoCas9, with 96,95 % of mutated loci with little influence on the promoter choice.

**Figure 2.**
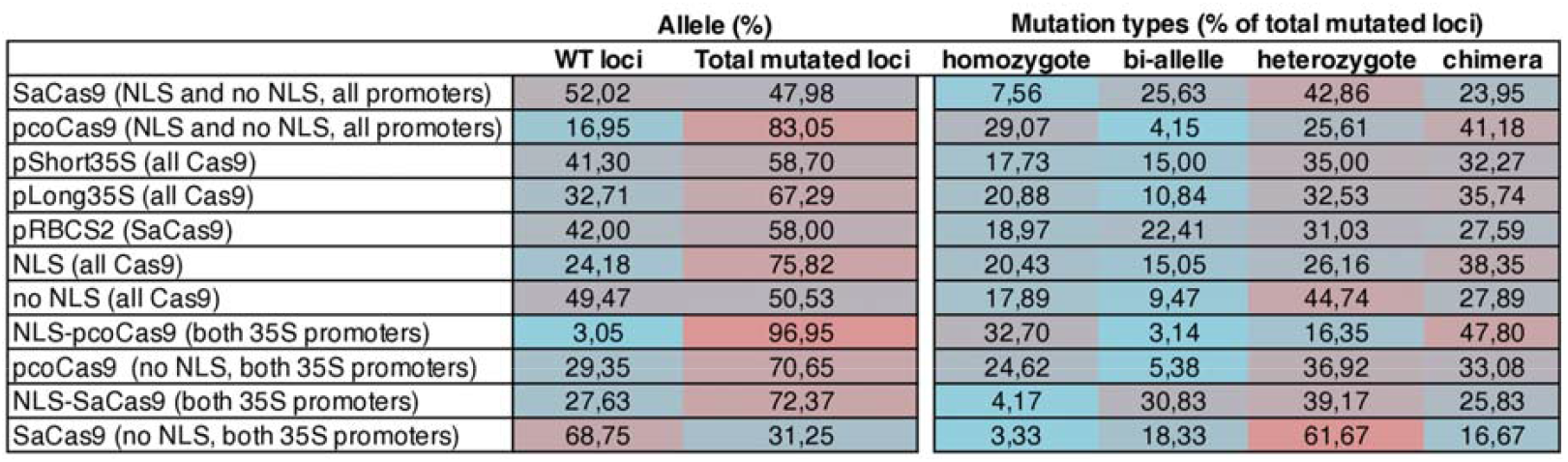
Table summarizing the mutagenesis efficiency of the CRISPR/Cas9 constructs for gRNA loci in *BnaA02*.*TAA1* (guide1 and guide2) and *BnaC02*.*TAA1* (guide1 and guide2). The mutagenesis efficiency is calculated as the percentage of the type of mutation on the total loci per construct (16 - 30 independent hairy root lines per construct). Data are pulled for a comparison of efficiency. The percentages of the type of alleles per loci (WT, mutated loci) and types of mutations (homozygote, bi-allele, heterozygote, chimera) over the total number of (mutated) loci are counted. Homozygote: the two alleles of a locus have an identical mutation. Bi-allele: the two alleles of a locus have different mutations. Heterozygote: only one allele is mutated. Chimera: more than two mutations per locus. Supporting data are shown in Supplemental Table S4.

Four categories of mutations were identified: homozygote mutation when both alleles have the same mutation, bi-allele mutation when the two alleles have a different mutation, heterozygote mutation when only one allele is mutated, and chimera when more than two types of mutations were identified at the analyzed locus. However, because each cell of a hairy root may have a different type of mutation, what we classified as heterozygote mutation may well be a mixture of homozygous and wild-type loci. The distribution of the mutation categories among the mutated loci is variable. Homozygote mutation is mainly observed with pcoCas9 (32,7 % with NLS-pcoCas9). All SaCas9 constructs preferentially generated heterozygote mutant loci (31,03 % for *RBCS2* promoter, and 39,17 % and 61,67 % for *35S* promoters, depending on the presence or absence of NLS, respectively). And in the most efficient constructs (NLS-pcoCas9), the majority of the loci are mutated as chimera alleles (47,8 %) (**Figure 2**).

Then, we looked at the size of the indels (insertion and deletion) induced by the activity of the Cas9 proteins. Because each locus may have more than two types of alleles (e.g., chimera), the distribution of the indel size among the constructs was calculated over the total number of alleles rather than loci. Most of the mutations were +1-base insertion for all constructs, ranging from 55,14 % of the mutated alleles for *pLong35S:NLS-pcoCas9* to 81,08 % of the mutated alleles for *pShort35S:NLS-SaCas9*. Nuclear pcoCas9 proteins were the most effective for deleting two or more bases, with 30,83 % of the mutated alleles for *pShort35S:NLS-pcoCas9* and 30,84 % for *pLong35S:NLS-pcoCas9* (**Figure 3A**).

**Figure 3.**
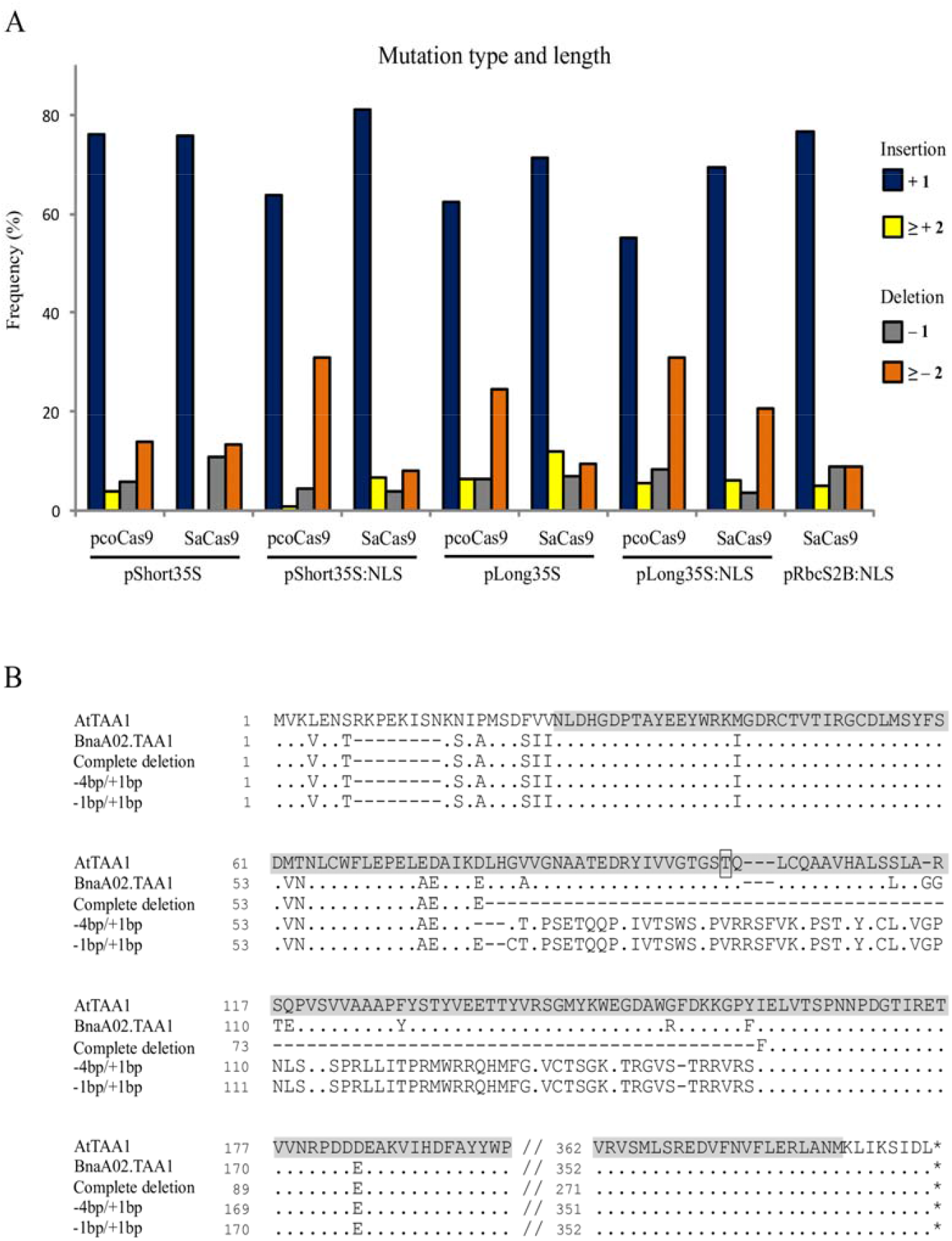
Size of the indels and effects of large deletions. A, Frequency of CRISPR/Cas9-induced mutations for each construct in relation to the size of the indel: insertion of one base (dark blue), insertion of 2 or more bases (yellow), deletion of 1 base (grey), and deletion of two or more bases (orange). The frequency is calculated as the number of mutated loci in the given category over the total mutated loci with the analyzed construct. B, Protein sequence alignment of TAA1. Comparison of *A. thaliana* ortholog (AtTAA1) with wild-type allele of BnaA02.TAA1 and deduced amino acids from CRISPR/Cas9 mutants with a sequence deletion between the two guides loci (complete deletion) or two specific combinations of mutations at exon II and exon III (−4-base or -1-base deletion combined with +1-base insertion) enabling ORF restoration. Dots and dashes represent identical and deleted nucleotides, respectively. The alliinase domain in AtTAA1 is highlighted in grey and the phosphorylation site at T101 is boxed.

We noticed that large deletions occurred between the two gRNA target loci in a few transformants. Complete deletion of the sequence between the two targeted loci, creating the loss of partial sequences of exon II and exon III, and the whole intron II, was identified three times for *BnaA02*.*TAA1* with pcoCas9 constructs. The coding sequence of *BnaA02*.*TAA1* was reduced to 903 bp instead of 1 146 bp, resulting in the deletion of 81 amino acids in the middle of the protein (**Figure 3B**).

Most of the indel for *BnaA02*.*TAA1* was a +1-base insertion in the 5’ targeted locus (exon II), leading to an ORF shift. A premature stop codon occurs 255 (or 267) bases downstream to the insertion depending on the presence (or not) of a second +1-base insertion at the second locus in exon III. In some transformants, the mutation in exon II (−1 or -4-base deletion) is compensated by +1-base insertion in exon III. In such a case, the protein sequence between the two loci is aberrant but restored after the second mutation (**Figure 3B**). The protein domain in *BnaA02*.*TAA1* affected by a large deletion or aberrant sequence between the two targeted loci is the alliinase domain, responsible for the aminotransferase activity of TAA1 protein. The phosphorylation site at Threonine 101 controlling the on–off switch of TAA1 enzymatic activity (Wang et al., 2020) is missing in these mutant proteins. Thus, such proteins may be valuable to study the functional activity of BnaTAA1 *in vivo*. In *BnaC02*.*TAA1*, a +1-base insertion at the 5’ targeted locus led to an ORF shift and a premature stop codon upstream to the second target locus. In such a case, the type of mutation at the second locus did not influence the resulted ORF sequence. An altered protein sequence resulted from a combined mutation in exon II and exon III (−9-base deletion / -3-base deletion) with missing three amino acids (IKEL>M) in position AA69 and one amino acid (FIE>FE) at AA149.

Considering the mutagenesis efficiency, the type, and size of the mutation, we concluded that nuclear targeted Cas9 proteins are the most efficient, with a preference for pcoCas9. The long *35S* promoter slightly improved the mutagenesis occurrence. We observed that both pcoCas9 and SaCas9 favor a +1-base insertion over the other indels, resulting in ORF shift and premature stop codon.

### Optimization of hairy roots regeneration and regeneration of CRISPR/Cas9 *BnaTAA1* mutants

The regeneration of plants is a highly variable process depending on the species and, even, ecotypes and cultivars. Thus, we developed a regeneration protocol for the *B. napus* DH12075 cultivar by modifying a previously published method for *Brassica* spp. (Christey and Sinclair, 1992). Increasing the auxin (1-Naphthaleneacetic acid, NAA) level to 8 mg/l in the regeneration medium enhanced almost twice the regeneration capacity of the DH12075 cultivar (**Supplemental Figure S3A**). The rooting of the shoots was improved by adding auxin (0,5 mg/l Indole-3-butyric acid, IBA) into the root induction medium (RIM) or by increasing the concentration of gelling agents (**Supplemental Figure S3B**). Only one of ten DH12075 wild-type hairy root lines did not regenerate. The optimized protocol (6-Benzylaminopurine, BAP 5 mg/l and NAA 8 mg/l in the regeneration medium, and 0,3 % phytagel with 0,5 mg/l IBA in RIM) was successfully used for regenerating plants from hairy root cultures of Topas, a variety referred to as being recalcitrant to transformation and plant regeneration.

Selected hairy root clones carrying a targeted mutation in *BnaTAA1* genes and non-mutated root lines were regenerated using the optimized protocol. In total, 125 T0 plants originating from 71 hairy root lines (1–2 regenerant(s) per line, all constructs included) were screened for the presence of the *Cas9* transgene and *Ri* T-DNA (TL and TR regions) in the plant genome. The *Cas9* transgene and TL-DNA were detected in genomic DNA extracted from leaves of all regenerants. Ninety-seven plants (77,6 %) carried the TR region of T-DNA. The absence of *Agrobacteria* contamination was confirmed by the absence of the *virC* locus in extracted DNA. The same genotyping profile was detected in DNA extracted from unpollinated pistils of selected plants.

Further, we studied the frequency of additional mutagenesis during the regeneration process in both pcoCas9 and SaCas9 lines. We focused on homozygous loci (either mutated or wild-type). In 2 out of 22 analyzed loci (9,1 %), new mutations were detected in plants regenerated from hairy root clones possessing pcoCas9 constructs. One of these new mutations was detected in a wild-type locus in the original hairy root clone. A similar result (13,6 %, 3/22) was observed for regenerants derived from hairy roots with SaCas9 cassettes with no new mutation detected in wild-type loci (**Supplemental Table S5**). The same (stable or new) mutations were detected in leaves and unpollinated pistils, suggesting that the additional mutagenesis event (+1-base insertion or short deletions ranging from -1 bp to -10 bp) occurred in the early stages of the regeneration process.

Regenerants exhibited an altered phenotype, including extensive root growth, curled leaves, and dwarfism (**Supplemental Figure S4**). Fertile flowers could set seeds, although the seed production was reduced compared to wild-type plants.

### Off-target mutagenesis is limited to homoeolog loci

Using the CRISPR-P v2.0 web tool, we identified the loci in the oilseed rape genome predicted to be most likely off-targets of the four gRNAs, and the top-ranking sites were selected for analysis (**Table 1**). We amplified regions surrounding the off-target sites from selected T0 plants showing the highest mutation rate in *BnaA02*.*TAA1* and *BnaC02*.*TAA1* genes (mutation detected in all four loci, preferentially homozygous mutations). For SaGuide1 and SaGuide2 analysis, genomic DNA of 15 edited plants (3 plants per each SaCas9 construct, i.e., *pShort35S:SaCas9, pLong35S:SaCas9, pShort35S:NLS-SaCas9, pLong35S:NLS-SaCas9, pRbcS2B:NLS-SaCas9*) was extracted from leaves, and potential off-target regions were amplified and sequenced. For pcoGuide1 and pcoGuide2 analysis, 12 edited plants (3 plants per each pcoCas9 construct, i.e., *pShort35S:pcoCas9, pLong35S:pcoCas9, pShort35S:NLS-pcoCas9, pLong35S:NLS-pcoCas9*) were tested.

The sequence of the amplified predicted off-target site for pcoGuide1 on chromosome A09 was divergent from the gRNA sequence. This discrepancy may result from genomic sequence differences between the cultivar used for transformation (DH12075) and the reference genome (Darmorbzh) used for *in silico* analysis. Therefore, the “real” off-score is not known but probably lower than the 0,554 off-score provided by CRISPR-P v2.0.

Besides the intergenic regions, where no mutations were found in putative off-target sites, two gene loci (*BnaAnng22030D* and *BnaC06g43720D*) were predicted for off-target editing by gRNAs of both Cas9. These genes represent paralogs of the *BnaTAA1* family (**Supplemental Figure S1**). As expected from a predicted low off-score, no mutations were detected in the SaGuide1 putative off-target site in *BnaAnng22030D* and *BnaC06g43720D* genes. The target sequence of SaGuide2 in these two paralogs contains only one SNP localized 9 bp upstream of the PAM sequence, and a high off-score was predicted for these loci. Indeed, we detected mutations in 5 and 3 of the 15 tested plants in *BnaAnng22030D* and *BnaC06g43720D*, respectively (**Table 1**). Both heterozygous and homozygous mutations (mostly +1-base insertion and one -4-base deletion) were identified. As for pcoGuide2 off-targets, we identified a heterozygous +1-base insertion in the *BnaC06g43720D* locus in one plant (out of 12 tested), although the computed off-score for this locus was slightly lower than for intergenic regions where no mutation was detected. Even though no off-target editing was predicted in *BnaAnng22030D* for pcoGuide2, we sequenced this region in all tested plants to complete the off-target study of paralogous genes. No mutation was detected.

### *BnaTAA1* transcripts analysis confirmed a down-regulation of *BnaTAA1* in the mutant regenerants

We monitored *BnaA02*.*TAA1* and *BnaC02*.*TAA1* expression by qPCR in unpollinated pistils of transgenic regenerant plants (**Figure 4**). As a control, we used a regenerant derived from wild-type hairy roots (no CRISPR/Cas9 construct was used for transformation). As expected, no transcript level reduction was detected in a plant regenerated from hairy roots containing the CRISPR/Cas9 transgene but no mutation in the *BnaTAA1* genes. The absence of additional mutations arising during the regeneration process of such a plant was verified by sequencing the *BnaTAA1* genes in pistils. However, the analysis of two plants regenerated from hairy root lines with an active CRISPR/Cas9 transgene (specific mutations are highlighted in **Figure 4A**) revealed a reduced total *BnaTAA1* expression to less than 10 % of the control level (6,3 % and 5 % for lines expressing pcoCas9 and SaCas9 constructs, respectively) (**Figure 4B**).

**Figure 4.**
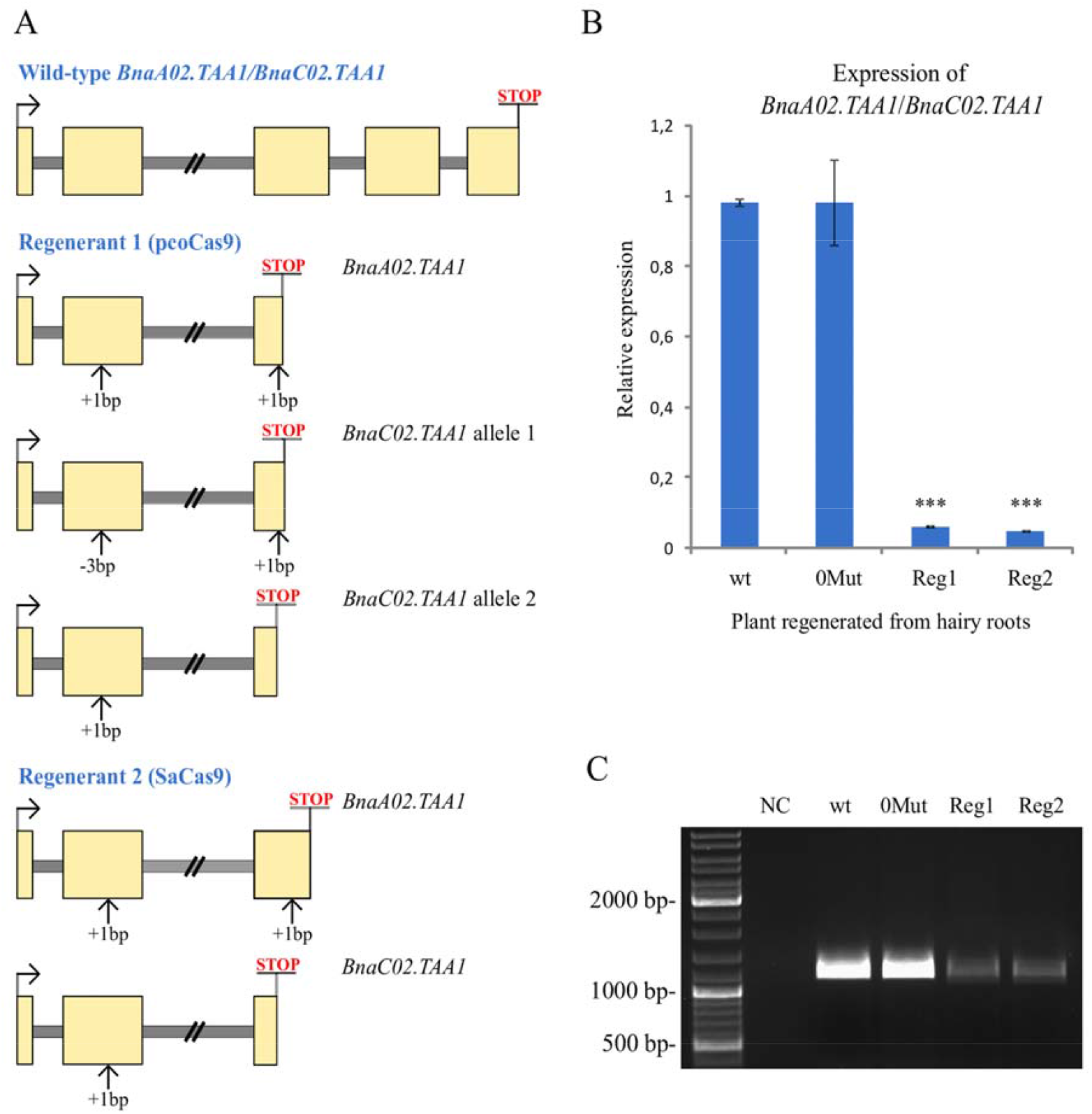
Expression analysis of *BnaTAA1* genes. A, Schematic representation of the indel influence on the coding regions in selected mutant lines. Exons are in yellow and introns in grey. The premature stop codon is highlighted. B, RT-qPCR analysis of *BnaTAA1* genes in selected T0 regenerated plants. wt, regenerant derived from wild-type hairy roots considered as a control. 0MUT, regenerant with no mutation in *BnaTAA1* genes. Reg1 and Reg2, *BnaTAA1* mutant plants. The data represent mean ± SD (n = 3) of normalized and rescaled expression levels. Significantly different expression levels are indicated by asterisks (one-way ANOVA; Dunnett’s Test; p < 0.0001). C, Amplification of the coding regions of *BnaA02*.*TAA1* and *BnaC02*.*TAA1* and partial 5’ and 3’ UTRs by RT-PCR. No alternative splicing of CRISPR/Cas9-targeted exons was detected in non-mutated regenerant (0MUT) or mutant plants (Reg1 and Reg2). NC, negative control of PCR.

In human cell lines, novel transcript variants have been detected in CRISPR/Cas9-edited cells. Alternative mRNA splicing was associated with targeted exons, as the indel-containing exon was excluded from mRNA in ∼30 % of the studied clones (Tuladhar et al., 2019). We employed the rapid amplification of cDNA ends (RACE) technique to determine the 5’ and 3’ UTRs of *BnaA02*.*TAA1* and *BnaC02*.*TAA1* genes. From these sequences, we designed primers to amplify the whole coding regions of *BnaA02*.*TAA1* and *BnaC02*.*TAA1* and partial 5’ and 3’ UTRs. No alternative splicing of targeted exons (exon II, 307 bp; and exon III, 290 bp) in CRISPR/Cas9-edited plants was detected (**Figure 4C**). The *BnaTAA1* expression level in the edited plants was consistent with the qPCR results.

### Effects of indels in *BnaTAA1* genes on T1 plants phenotypes

The *rol* genes encoded on the T-DNA of the *Ri* plasmid are integrated into the plant genome and are crucial for hairy-root formation. However, their presence in regenerated plants correlates with altered growth characteristics called Ri phenotype (**Supplemental Figure S4**). Another aspect that must be considered is the presence of the Cas9 transgene in the genome. Obtaining modified plants free of the Cas9 transgene ensures no additional genome editing. The insertion of the *Ri* T-DNA and the Cas9 transgenes may be independent or linked. Therefore, a segregation analysis of T1 progeny identified T-DNA-free mutant plants.

T0 plants from selected transgenic lines were selfed to produce seeds. At first, we screened the roots of T1 seedlings for the absence of a fluorescence signal, as all CRISPR/Cas9 constructs contain a GFP reporter. The absence of Cas9 transgene in GFP-negative seedlings was verified by PCR with primers designed to amplify the *Cas9* gene (**Supplemental Figure S4, Supplemental Table S1**). The Cas9-free plants were further genotyped for the absence of TL- and TR-DNA of the *Ri* plasmid. Cas9-free regenerants were subjected to *BnaTAA1* analysis. No additional mutations were detected in T1 progeny compared to the T0 plants (35 loci tested).

In Arabidopsis, disruption of *TAA1* alone does not cause dramatic developmental phenotypes, but inactivation of *TAA1* and *TAR2* homolog leads to defects in root and flower development (Stepanova et al., 2008). Homozygous single nucleotide insertion in *BnaC02*.*TAA1* gene and homozygous large deletion in *BnaA02*.*TAA1* in T1 mutant plants (**Figure 5A**) led to the *BnaTAA1* transcripts level reduction as observed by RT-PCR (**Figure 5B**). We observed a more compacted inflorescence in the main stem with more floral buds in mutant T1 plants. A few of those buds do not mature and senesce, which was not observed in the main wild-type inflorescence (**Figure 5C)**. The mutant plants have less developed siliques, often shorter and thinner than the wild-type plants (**Figure 5D**).

**Figure 5.**
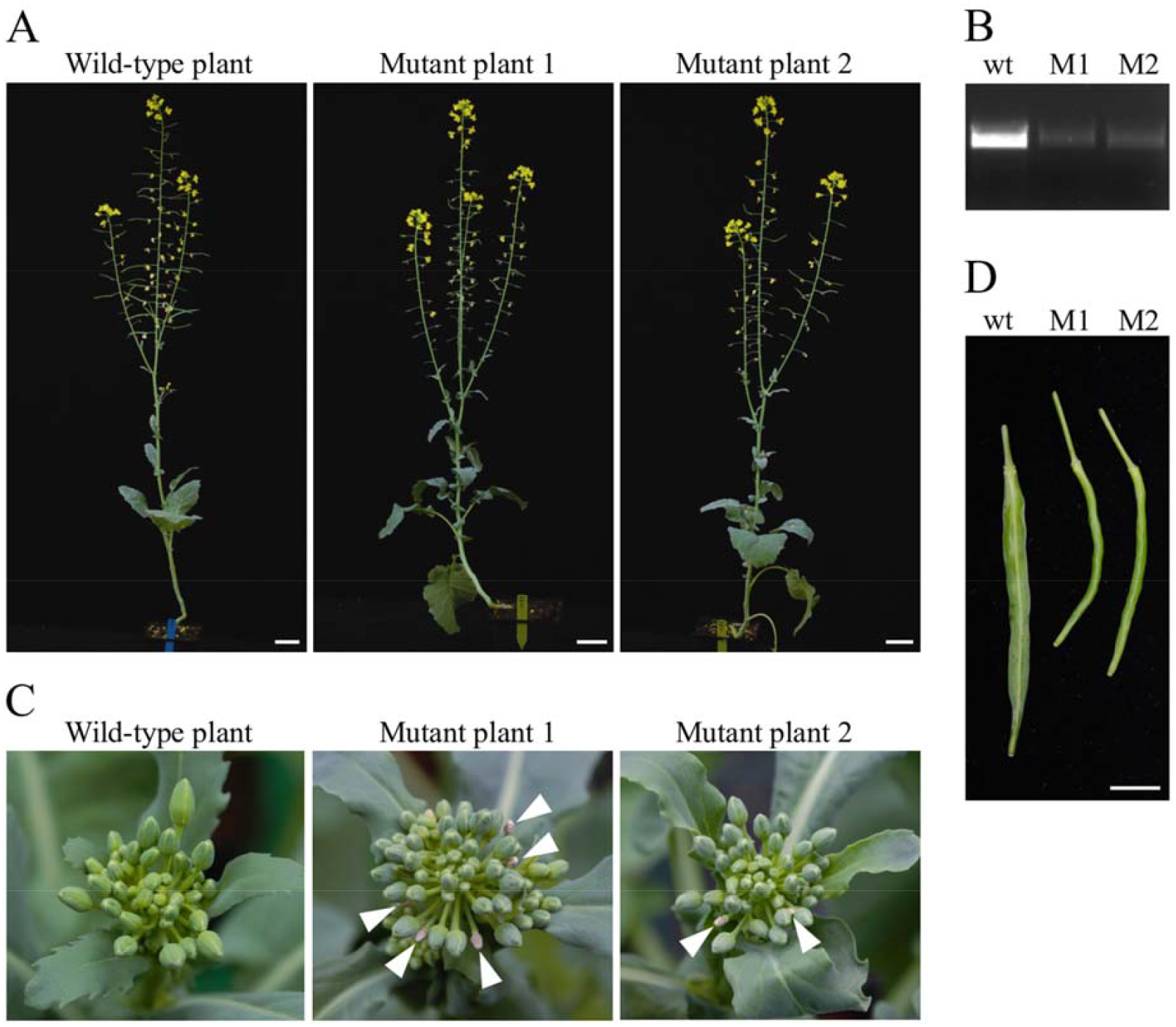
Phenotypes of *BnaTAA1* double mutants. A, Morphology of T1 plants, free of *Ri* T-DNA and Cas9 transgene, carrying the homozygous mutations in *BnaA02*.*TAA1* and *BnaC02*.*TAA1* at the same age as the wild-type *B. napus* DH12075 (scale bar = 5 cm). B, Homozygous indels lead to a reduced *BnaTAA1* expression detected by RT-PCR. C, Inflorescence meristem (top view) of mutants compared to the wild-type plant. Arrowheads indicate senescent flower buds. D, Mature green siliques of mutants compared to the wild-type plant (scale bar = 1 cm).

## Discussion

The hairy root transformation system as a tool for CRISPR/Cas9-based genome editing has been established recently for various species (reviewed in Kiryushkin et al., 2022). Among *Brassica* crops, *B. carinata* hairy roots were successfully edited using a CRISPR/Cas9 system (Kirchner et al., 2017). As *B. napus* is a tetraploid species, mutating homoeologs from both subgenomes is necessary due to functional genetic redundancy. Thus, an optimization of CRISPR/Cas9 constructs is essential, as the gene-editing efficiency depends on the selection of the Cas9 protein, the promoter driving the expression of the *Cas9* gene, or the selection of guide RNAs (Steinert et al., 2015; Bortesi et al., 2016). Hairy root transformation represents a fast and straightforward system for such evaluation, as, within approximately two months after transformation, it is possible to evaluate the most effective gene-editing construct (**Supplemental Figure S4**).

In our study, we performed the bioinformatic analysis of the amplified and sequenced targeted loci using TIDE (tracking of indels by decomposition) combined with the cloning of fragments with complex mutations to evaluate the frequencies and types of different mutations accurately. Alternatively, other methods to distinguish between wild-type and mutated loci can be used, such as restriction fragment length polymorphism (RFLP) technique, high resolution melting analysis (HRMA), or T7EI assays based on nuclease cleaving heteroduplex DNA fragments (Lomov et al., 2019).

We showed that the CRISPR/Cas9 constructs simultaneously mutated paralogous genes within the *BnaTAA1* gene family. We detected mutations in *BnaAnng22030D* and *BnaC06g43720D* paralogs in a significant portion of plants using a gRNA with one SNP in a target sequence. These genes are expressed at a very low level in oilseed rape plants (*Brassica* Expression Database, Chao et al., 2020), and their function can be investigated in plants with *BnaA02*.*TAA1/BnaC02*.*TAA1* loss-of-function mutants. As various combinations of mutated paralogs within the *BnaTAA1* gene family were achieved, the quantitative involvement of *BnaTAA1* copies in the auxin biosynthesis can be studied in *B. napus*.

Using a pair of gRNAs encoded in a single CRISPR array, two DSBs flanking the target DNA may result in the excision of the genomic DNA fragment. Such large deletion can lead to either a frameshift in coding sequence and premature stop codon or the ORF restoration forming a chimeric transcript variant. Transcripts carrying premature stop codon are usually degraded by nonsense-mediated mRNA decay (NMD) pathway (Lykke-Andersen and Jensen, 2015), as we detected for the transcripts with single bp mutation where the transcript abundance dropped under 10% compared to the control. Deletion preserving the reading frame may yield a protein with altered amino acid composition and shed light on the biological function of such protein. In our design, the deletion between two gRNAs target sites encompasses the phosphorylation site at Threonine 101 in the alliinase domain controlling the TAA1 enzymatic activity (Wang et al., 2020). We achieved such mutations with pcoCas9 constructs showing higher efficiency in deletion of ≥ 2 bases compared to the SaCas9 vectors. Among pcoCas9 constructs, pcoCas9-NLS induced homozygous mutations with the highest efficiency (∼ 30% of the mutated loci).

The CRISPR/Cas9 constructs with the highest efficiency of targeted mutagenesis can be used for traditional *A. tumefaciens*-based transformation of explants followed by regeneration of transgenic plants. Alternatively, the so-called composite plants consisting of wild-type shoots and transgenic hairy roots can be generated. This technique works efficiently for many plant species, including those recalcitrant to transformation (Taylor et al., 2006). The system allows for “in root” study of gene disruption in the context of a whole plant, similar to studies with grafted plants. The injection-based method used for *B. napus* transformation is an excellent choice for such studies as fully developed hairy roots emerging from the inoculation site can sustain plant growth after removing the original roots.

Another option is the regeneration of the existing hairy root lines with defined and desired mutations. We optimized the hairy root regeneration protocol for *B. napus* cultivar DH12075. In some hairy root lines, the shoots appeared within the first month of cultivation on regeneration media. In other lines, it took 2 – 3 months to regenerate shoots. Shoots were cultivated on shoot elongation and root induction media, and the rooted regenerated plants were subsequently transferred to the soil and cultivated for 3 – 5 months to produce immature seeds. An embryo rescue of 21 – 28 day-after-pollination seeds containing torpedo stage embryos or older was performed to germinate T1 plantlets, saving otherwise necessary weeks for seed maturation and dormancy. In total, it is possible to obtain transgene-free T1 plants with desired mutations roughly one year after agrobacterial transformation (**Supplemental Figure S4**).

The *taa1* single mutants in Arabidopsis display mild auxin-deficient phenotypes (Stepanova et al., 2008) that correspond to our observations in *B. napus*. We detected flower and silique development defects in plants with reduced *BnaA02*.*TAA1/BnaC02*.*TAA1* transcripts level caused by CRISPR/Cas9-induced mutations. A set of plants with various mutations in the *BnaTAA1* family will serve as a tool for understanding genotype-phenotype correlations.

## Conclusions

Using hairy root transformation, we successfully applied CRISPR/Cas9 system to generate and evaluate gene-editing in the *BnaTAA1* target genes in *B. napus*. We demonstrated that gRNAs designed on conserved sequences could simultaneously induce mutations at multiple loci, which are stable and heritable from hairy roots to regenerated plants. Constructs expressing two gRNAs for targeted genes enabled both single base-pair indels and deletion of the whole sequence between the two target sites.

## Materials and Methods

### Plant material

For the transformation experiments, seeds of three *Brassica napus* cultivars (DH12075, Westar, and Topas DH4079) were sterilized with 20 % bleach and cultivated *in vitro* on 50 % Murashige and Skoog (MS) medium (Duchefa) with 5 g/L sucrose at 21 °C (16 h photoperiod) in a growth chamber. Plants growing in the soil were cultivated in a greenhouse with similar temperature and light conditions.

### *BnaAnng22030D* cloning and phylogenetic analysis

The genomic sequence corresponding to the *BnaAnng22030D* gene and its surroundings was amplified using specific primers (**Supplemental Table S1**) and PrimeSTAR GXL DNA Polymerase (Takara). The PCR product was cloned and sequenced. Its sequence was deposited in the GenBank database (OM687491).

Multiple alignments of protein sequences were performed using Muscle software (Edgar, 2004), and phylogenetic analysis was carried out using the neighbor-joining method with the SeaView program (Gouy et al., 2010). Bootstrap values were calculated from 1000 replications. The resulting phylogenetic tree was drawn and edited using FigTree v.1.4.4 (http://tree.bio.ed.ac.uk/software/figtree/).

### CRISPR/Cas9 vector construction

The two guide RNAs (gRNA) targeting the coding sequences of *BnaTAA1* genes *BnaC02g19980D* and *BnaA02g14990D* were designed for two Cas9 nucleases (*Staphylococcus aureus* SaCas9 and plant codon-optimized *Streptococcus pyogenes* Cas9, pcoCas9) using CRISPR-P v2.0 online tool (http://crispr.hzau.edu.cn/CRISPR2/) (Lei et al., 2014; Ding et al., 2016; Liu et al., 2017). For plasmid construction, a modular cloning system was employed using MoClo Tool Kit and MoClo Plant Parts kit (Addgene, **Supplemental Table S3)** (Weber et al., 2011; Werner et al., 2012; Engler et al., 2014). Plasmids were assembled according to the recommended long MoClo protocol (http://synbio.tsl.ac.uk/). Eight combinations of long (1,3 kb) and short (0,4 kb) versions of the *35S* promoter from the Cauliflower mosaic virus and the two Cas9 nucleases were generated (**Figure 1**). A ninth construct containing the Arabidopsis *RbcS2B* promoter driving the expression of *SaCas9* was included. Each plasmid comprises a plant kanamycin resistance gene, a gene expressing GFP to facilitate the screening process, and the two respective gRNA cassettes (**Figure 1**). The intermediate constructs for gRNA were prepared as universal plasmids containing an AtU6-26 promoter, followed by two Esp3I sites for gRNA insertion instead of a counter-selective LacZ gene and a scaffold gRNA sequence. Specific guides were synthesized as oligonucleotides with added Esp3I sites, which enable scarless insertion, annealed by boiling and slow cooling, and used for standard assembly protocol with the prepared universal plasmids (details in **Supplemental Table S2**). Plasmids were sequenced before their use.

### Plant transformation (hairy root cultures)

Hairy root cultures were obtained by transforming *B. napus* plants with the *Ti*-less plasmid *Agrobacterium tumefaciens* C58C1 strain carrying a hairy-root-inducing plasmid *pRiA4b* (Petit et al., 1983). The transformation assay was optimized with *Agrobacterium* containing only the *Ri* plasmid (root lines are designated as wild-type hairy roots). For CRISPR/Cas9 analysis, each construct was electroporated into *Agrobacterium*. A selected bacterial clone was verified by PCR analysis with construct-specific primers (**Supplemental Table S1**). For plant transformation, an *Agrobacterium* suspension was grown for 18 hours in Luria-Broth medium at 28 °C and injected with an insulin syringe into the hypocotyl of 18-day-old seedlings cultivated *in vitro*. After 2 – 4 cultivation weeks, hairy roots emerging from the inoculation sites were excised and placed on a solid MS medium, including Gamborg B5 vitamins (MS+B5, Duchefa) and 30 g/L sucrose, supplemented with ticarcillin (500 mg/l) and cefotaxime (200 mg/l) to eliminate bacteria growth. In case of transformation with a CRISPR/Cas9 construct, kanamycin (25 mg/L) was added to the medium. Hairy roots were grown in Petri dishes at 24 °C in the dark and transferred to fresh MS+B5 media after 4 – 5 cultivation weeks. The concentration of ticarcillin and cefotaxime was decreased during each transfer. After 3 – 4 months, the roots were maintained on solid MS containing cefotaxime (100 mg/L) only. To support the growth of mutated lines in the auxin biosynthesis *BnaTAA1* genes, we added a low concentration (0,25 mg/L) of indole-3-butyric acid (IBA) into the media.

### Genomic DNA extraction (hairy roots, leaves, pistils)

Hairy roots were collected from Petri dishes, and remnants of solid media were carefully removed. Young leaves were sampled before the flowering stem emerged. For unpollinated pistils, the oldest flower buds were emasculated one day before the flower opening. On the next day, the pistils were sampled. Tissues were ground in liquid nitrogen, and genomic DNA was isolated by the conventional cetyltrimethylammonium bromide (CTAB) method (Allen et al., 2006).

### Analysis of gene editing in hairy roots

One to three roots per independent transformant and 13 – 20 transformants per construct (16 – 30 transformed hairy roots per construct) were analyzed. The *BnaTAA1* genes were amplified by PCR to analyze the mutations mediated by CRISPR/Cas9, using Phusion High-Fidelity DNA polymerase (New England Biolabs) and gene-specific primers discriminating between *BnaA02*.*TAA1* and *BnaC02*.*TAA1* (**Supplemental Table S1**), and sequenced. The online tool TIDE (https://tide.nki.nl/) was used for chromatogram decoding (Brinkman et al., 2014). The TIDE results were confirmed manually. For the more complex mutations, amplified fragments were subcloned into the pGEM-T-Easy Vector (Promega), and 6 – 10 clones of each amplicon were sequenced. The type and length of the indels (insertion/deletion) were recorded.

### Regeneration of hairy root clones

Optimization of hairy root regeneration was carried out by transferring ten independent wild-type hairy root clones to MS+B5 solid media with 30 g/L sucrose supplemented with cytokinin 6-benzylaminopurine (BAP, 5 mg/L) and various concentrations of auxin, 1-naphthaleneacetic acid (NAA, 0 – 8 mg/L), and indole-3-butyric acid (IBA, 0 – 8 mg/L). Roots were grown at 21 °C (16 h photoperiod) and transferred to fresh media every 3 – 4 weeks. The evaluation of the regeneration efficiency was performed after 3 months (**Supplemental Figure S3**). The excised shoots were cultivated on a shoot elongation medium (MS+B5 with 20 g/L sucrose supplemented with 0,5 mg/L BAP and 0,03 mg/L gibberellic acid). After 3 – 4 weeks, they were transferred to the root induction medium (RIM; MS+B5 with 10 g/L sucrose). The influence of the gelling agent (phytagel) concentration in RIM and the addition of IBA were studied (**Supplemental Figure S3**). Rooted plants were transferred to the soil. The optimized protocol (BAP 5 mg/l and NAA 8 mg/l in regeneration medium and 0,3% phytagel with 0,5 mg/l IBA in RIM) was used to regenerate CRISPR/Cas9 transformants. Genomic DNA isolated from T0 regenerant leaves was used for mutation genotyping and detection of CRISPR/Cas9 transgene, *Ri* T-DNA (TL and TR), and *virC* genes (oligo information in **Supplemental Table S1**).

### Embryo rescue, T1 seedlings screening, plant phenotyping

To speed up the workflow with the selection of T1 plants, immature T1 seeds (21 – 28 day-after-pollination) containing torpedo stage embryos or older were embryo rescued, bypassing the maturation and dormancy phase. Siliques were collected and surface-sterilized with 70% ethanol. Under sterile conditions, the siliques were slit-opened, and the immature seeds were carefully collected without damaging the seed coat. The seeds were germinated on 50 % MS plates in the growth chamber. Seedlings were transferred to a hydroponic culture system to allow root tip fluorescence screening. Genomic DNA isolated from T1 GFP-negative seedlings was used to detect CRISPR/Cas9 transgene and *Ri* T-DNA (TL and TR). The transgene-free plants were used for mutation genotyping (*BnaA02*.*TAA1* and *BnaC02*.*TAA1* were screened for one or both loci; 35 loci in total) (oligo information in **Supplemental Table S1**). T1 transgene-free mutant plants were grown in soil side-by-side with DH12075 cultivars in the greenhouse at 21 °C (16 h photoperiod), flowers were self-pollinated, and the plants were monitored for phenotypes

### RNA analysis

Total RNA was extracted from unpollinated pistils using TRIzol reagent (Invitrogen) following the manufacturer’s protocol. RNA isolates were treated with RQ1 RNase-Free DNase (Promega) to remove traces of contaminant DNA. To obtain the 3’ ends of the *BnaTAA1* transcripts, we performed reverse transcription using M-MLV Reverse Transcriptase (Promega) on 2 μg of template RNA with an oligo-dT primer. The 5’ ends of *BnaTAA1* transcripts were detected using FirstChoice RLM-RACE Kit (Invitrogen) with 5 μg of the template RNA according to the manufacturer’s instructions. Primers used for subsequent cDNA amplification are listed in **Supplemental Table S1**. PCR products were cloned and sequenced. The cDNA sequences of *BnaA02*.*TAA1* and *BnaC02*.*TAA1* genes are available from GenBank under accession numbers OM687489 and OM687490, respectively. Primers recognizing 5’UTR and 3’UTR of both *BnaTAA1* genes (**Supplemental Table S1**) were used to study alternative splicing.

For RT-qPCR analysis, cDNA synthesis was performed on 1,5 μg of RNA using M-MLV Reverse Transcriptase (Promega). PCR reaction was performed using the FastStart Essential DNA Green Master (Roche) on a Lightcycler 96 (Roche). The efficiency of each primer pair (**Supplemental Table S1**) was assessed by constructing a standard curve through five serial dilutions. A final meltcurve step was included post-PCR to confirm the absence of any non-specific amplification. The experiment consisted of three independent biological replicates with three technical replicates for each parallel group. The suitability of reference genes (ACT7 [LOC106426760, LOC106384924, LOC106441419]; EF1A [LOC111197859, LOC106400771, LOC106419721, LOC106385889], and TBP2 [LOC106361251, LOC106360359]) was evaluated using mathematical methods implemented in BestKeeper (Pfaffl et al., 2004), geNorm (Vandesompele et al., 2002), and NormFinder (Andersen et al., 2004). TATA-box-binding protein 2 (TBP2), the most reliable reference gene, was used in the subsequent analyses. Relative gene expression was determined using the method described by Pfaffl (2001). The expression levels were evaluated by one-way ANOVA followed by Dunnett’s Test.

### Off-target analysis

The potential off-target sites were predicted using CRISPR-P v2.0 online tool. The top-ranking sites were selected for analysis. The genomic DNA sequences surrounding the potential off-target sites were amplified by PCR using specific primers (**Supplemental Table S1**) and PrimeSTAR GXL DNA Polymerase (Takara). PCR products were analyzed by sequencing.

### Accession numbers

*BnaA02*.*TAA1* cDNA OM687489; *BnaC02*.*TAA1* cDNA OM687490; *BnaAnng22030D* genomic sequence OM687491.

## Acknowledgments

The authors acknowledge Kim Boutilier (Wageningen University, The Netherlands) for providing the *Brassica napus* seeds, Jiří Macas (Biology Centre CAS, České Budějovice, CR) for providing the agrobacterial strain, Ben Miller (University of East Anglia, Norwich, UK) for providing the pcoCas9 plasmids, and Holger Puchta (Karlsruher Institute for Technology, Germany) for providing the SaCas9 plasmids. The Moclo Plant Parts kit was a gift from Nicola Patron (Addgene kit #1000000047). The Moclo Toolkit was a gift from Sylvestre Marillonnet (Addgene kit #1000000044). The authors acknowledge the core facility CELLIM supported by the Czech-BioImaging large RI project (LM2018129 funded by MEYS CR) for their support in obtaining scientific data presented in this paper. The Plant Sciences Core Facility of CEITEC Masaryk University is acknowledged for its technical support.

## Author contributions

V.J. designed and performed the hairy root culture experiment and CRISPR-mutagenesis screen, analyzed and interpreted the data, wrote the manuscript draft. J.Z. and K.M. designed and cloned the CRISPR plasmids. K.M. screened and sequenced the mutated sites. J.B. optimized the hairy root protocol and plant regeneration. M.St. monitored the plants (growth, picture), performed the RT-qPCR expression analysis. M.Se. genotyped the plants. H.S.R. conceived the project, analyzed the CRISPR-induced mutations, wrote and reviewed the manuscript. H.S.R. agrees to serve as the author responsible for contact and ensures communication. All authors have read and approved the final version of the manuscript.

## Supplemental Material

**Supplemental Table S1**. List of oligonucleotides

**Supplemental Table S2**. Plasmid construction for CRISPR/Cas9-mediated genome editing using MoClo system

**Supplemental Table S3**. Origin of plasmids

**Supplemental Table S4**. Mutagenesis efficiency of each construct for each gRNA loci

**Supplemental Table S5**. Stability of homozygous loci in hairy root regenerants

**Supplemental Figure S1**. Phylogenetic analysis of TAA1, TAR1, and TAR2 proteins in *Brassica napus* and related species

**Supplemental Figure S2**. Hairy roots induction in different *B. napus* cultivars

**Supplemental Figure S3**. Optimization of hairy roots regeneration in *B. napus* DH12075

**Supplemental Figure S4**. Workflow scheme for transgene-free *BnaTAA1*-edited plants generation

## Parsed Citations

Allen GC, Flores-Vergara MA, Krasynanski S, Kumar S, Thompson WF (2006) A modified protocol for rapid DNA isolation from plant tissues using cetyltrimethylammonium bromide. Nat Protoc 1: 2320–2325

Agostini E, Talano MA, González PS, Oller AL, Medina MI (2013) Application of hairy roots for phytoremediation: what makes them an interesting tool for this purpose? Appl Microbiol Biotechnol 97: 1017–1030

Andersen CL, Jensen JL, Ørntoft TF (2004) Normalization of real-time quantitative reverse transcription-PCR data: a model-based variance estimation approach to identify genes suited for normalization, applied to bladder and colon cancer data sets. Cancer Res 64: 5245–5250

Bortesi L, Zhu C, Zischewski J, Perez L, Bassié L, Nadi R, Forni G, Lade SB, Soto E, Jin X, Medina V, Villorbina G, Muñoz P, Farré G, Fischer R, Twyman RM, Capell T, Christou P, Schillberg S (2016) Patterns of CRISPR/Cas9 activity in plants, animals and microbes. Plant Biotechnol J 14: 2203–2216

Brinkman EK, Chen T, Amendola M, van Steensel B (2014) Easy quantitative assessment of genome editing by sequence trace decomposition. Nucleic Acids Res 42: e168–e168

Cardarelli M, Mariotti D, Pomponi M, Spanò L, Capone I, Costantino P (1987) Agrobacterium rhizogenes T-DNAgenes capable of inducing hairy root phenotype. Mol Gen Genet 209: 475–480

Chao H, Li T, Luo C, Huang H, Ruan Y, Li X, Niu Y, Fan Y, Sun W, Zhang K, Li J, Qu C, Lu K (2020) BrassicaEDB: AGene Expression Database for Brassica Crops. Int J Mol Sci 2: 5831

Christey MC (2001) Use of ri-mediated transformation for production of transgenic plants. In Vitro Cell Dev Biol Plant 37: 687–700

Christey MC, Sinclair BK (1992) Regeneration of transgenic kale (Brassica oleracea var. acephala), rape (B. napus) and turnip (B. campestris var. rapifera) plants via Agrobacterium rhizogenes mediated transformation. Plant Sci 87: 161–169

Deltcheva E, Chylinski K, Sharma CM, Gonzales K, Chao Y, Pirzada ZA, Eckert MR, Vogel J, Charpentier E (2011) CRISPR RNA maturation by trans-encoded small RNAand host factor RNase III. Nature 471: 602–607

Ding Y, Li H, Chen L-L, Xie K (2016) Recent Advances in Genome Editing Using CRISPR/Cas9. Front Plant Sci 7: 703

Doudna JA, Charpentier E (2014) Genome editing. The new frontier of genome engineering with CRISPR-Cas9. Science 346: 1258096

Edgar RC (2004) MUSCLE: multiple sequence alignment with high accuracy and high throughput. Nucleic Acids Res 32: 1792–1797

Engler C, Youles M, Gruetzner R, Ehnert T-M, Werner S, Jones JDG, Patron NJ, Marillonnet S (2014) Agolden gate modular cloning toolbox for plants. ACS Synth Biol 3: 839–843

Gelvin SB (1990) Crown gall disease and hairy root disease: a sledgehammer and a tackhammer. Plant Physiol 92: 281–285

Gelvin SB (2003) Agrobacterium-mediated plant transformation: the biology behind the “gene-jockeying” tool. Microbiol Mol Biol Rev 67: 16–37

Georgiev MI, Agostini E, Ludwig-Müller J, Xu J (2012) Genetically transformed roots: from plant disease to biotechnological resource. Trends Biotechnol 30: 528–537

Gocal GFW (2021) Gene editing in Brassica napus for basic research and trait development. In Vitro Cell Dev Biol Plant 57: 731–748

Gouy M, Guindon S, Gascuel O (2010) SeaView version 4: a multiplatform graphical user interface for sequence alignment and phylogenetic tree building. Mol Biol Evol 27: 221–224

Grützner R, Martin P, Horn C, Mortensen S, Cram EJ, Lee-Parsons CWT, Stuttmann J, Marillonnet S (2021) High-efficiency genome editing in plants mediated by a Cas9 gene containing multiple introns. Plant Commun 2: 100135

Gutierrez-Valdes N, Häkkinen ST, Lemasson C, Guillet M, Oksman-Caldentey KM, Ritala A, Cardon F (2020) Hairy Root Cultures-A Versatile Tool With Multiple Applications. Front Plant Sci 11: 33

Jinek M, Chylinski K, Fonfara I, Hauer M, Doudna JA, Charpentier E (2012) Aprogrammable dual-RNA-guided DNAendonuclease in adaptive bacterial immunity. Science 337: 816–821

Kaya H, Mikami M, Endo A, Endo M, Toki S (2016) Highly specific targeted mutagenesis in plants using Staphylococcus aureus Cas9. Sci Rep 6: 26871

Keller O, Kollmar M, Stanke M, Waack S (2011) Anovel hybrid gene prediction method employing protein multiple sequence alignments. Bioinformatics 27: 757–763

Kirchner TW, Niehaus M, Debener T, Schenk MK, Herde M (2017) Efficient generation of mutations mediated by CRISPR/Cas9 in the hairy root transformation system of Brassica carinata. PLoS One 12: e0185429

Kiryushkin AS, Ilina EL, Guseva ED, Pawlowski K, Demchenko KN (2021) Hairy CRISPR: Genome Editing in Plants Using Hairy Root Transformation. Plants (Basel) 11: 51

Lei Y, Lu L, Liu H-Y, Li S, Xing F, Chen L-L (2014) CRISPR-P: A Web Tool for Synthetic Single-Guide RNADesign of CRISPR-System in Plants. Mol Plant 7: 1494–1496

Li J-F, Norville JE, Aach J, McCormack M, Zhang D, Bush J, Church GM, Sheen J (2013) Multiplex and homologous recombination–mediated genome editing in Arabidopsis and Nicotiana benthamiana using guide RNAand Cas9. Nat Biotechnol 31: 688–691

Li X, Sandgrind S, Moss O, Guan R, Ivarson E, Wang ES, Kanagarajan S, Zhu LH (2021) Efficient Protoplast Regeneration Protocol and CRISPR/Cas9-Mediated Editing of Glucosinolate Transporter (GTR) Genes in Rapeseed (Brassica napus L.). Front Plant Sci 12: 680859

Lin CS, Hsu CT, Yang LH, Lee LY, Fu JY, Cheng QW, Wu FH, Hsiao HC, Zhang Y, Zhang R, Chang WJ, Yu CT, Wang W, Liao LJ, Gelvin SB, Shih MC (2018) Application of protoplast technology to CRISPR/Cas9 mutagenesis: from single-cell mutation detection to mutant plant regeneration. Plant Biotechnol J 16: 1295–1310

Liu H, Ding Y, Zhou Y, Jin W, Xie K, Chen L-L (2017) CRISPR-P 2.0: An Improved CRISPR-Cas9 Tool for Genome Editing in Plants. Mol Plant 10: 530–532

Lykke-Andersen S, Jensen TH (2015) Nonsense-mediated mRNA decay: an intricate machinery that shapes transcriptomes. Nat Rev Mol Cell Biol 16: 665–677

Lomov NA, Viushkov VS, Petrenko AP, Syrkina MS, Rubtsov MA(2019) Methods of Evaluating the Efficiency of CRISPR/Cas Genome Editing. Mol Biol 53: 862–875

Mao Y, Zhang Z, Feng Z, Wei P, Zhang H, Botella JR, Zhu JK (2016) Development of germ-line-specific CRISPR-Cas9 systems to improve the production of heritable gene modifications in Arabidopsis. Plant Biotechnol J 14: 519–532

Petit A, David C, Dahl GA, Ellis JG, Guyon P, Casse-Delbart F, Tempé J (1983) Further extension of the opine concept: Plasmids in Agrobacterium rhizogenes cooperate for opine degradation. Mol Gen Genet 190: 204–214

Pfaffl MW (2001) Anew mathematical model for relative quantification in real-time RT-PCR. Nucleic Acids Res 29: e45

Pfaffl MW, Tichopad A, Prgomet C, Neuvians TP (2004) Determination of stable housekeeping genes, differentially regulated target genes and sample integrity: BestKeeper--Excel-based tool using pair-wise correlations. Biotechnol Lett 26: 509–515

Steinert J, Schiml S, Fauser F, Puchta H (2015) Highly efficient heritable plant genome engineering using Cas9 orthologues from Streptococcus thermophilus and Staphylococcus aureus. Plant J 84: 1295–1305

Stepanova AN, Robertson-Hoyt J, Yun J, Benavente LM, Xie DY, Dolezal K, Schlereth A, Jürgens G, Alonso JM (2008) TAA1-mediated auxin biosynthesis is essential for hormone crosstalk and plant development. Cell 133: 177–191

Taylor CG, Fuchs B, Collier R, Lutke WK (2006) Generation of composite plants using Agrobacterium rhizogenes. Methods Mol Biol 343: 155–167

Tuladhar R, Yeu Y, Tyler Piazza J, Tan Z, Rene Clemenceau J, Wu X, Barrett Q, Herbert J, Mathews DH, Kim J, Hyun Hwang T, Lum L (2019) CRISPR-Cas9-based mutagenesis frequently provokes on-target mRNA misregulation. Nat Commun 10: 4056

Vandesompele J, Preter KD, Pattyn F, Poppe B, Roy NV, Paepe AD, Speleman F (2002) Accurate normalization of real-time quantitative RT-PCR data by geometric averaging of multiple internal control genes. Genome Biol 3: RESEARCH0034

Wan H, Cui Y, Ding Y, Mei J, Dong H, Zhang W, Wu S, Liang Y, Zhang C, Li J, Xiong Q, Qian W (2017) Time-Series Analyses of Transcriptomes and Proteomes Reveal Molecular Networks Underlying Oil Accumulation in Canola. Front Plant Sci 7: 2007

Wang Q, Qin G, Cao M, Chen R, He Y, Yang L, Zeng Z, Yu Y, Gu Y, Xing W, Tao WA, Xu T (2020) Aphosphorylation-based switch controls TAA1-mediated auxin biosynthesis in plants. Nat Commun 11: 679

Wang ZP, Xing HL, Dong L, Zhang HY, Han CY, Wang XC, Chen QJ (2015) Egg cell-specific promoter-controlled CRISPR/Cas9 efficiently generates homozygous mutants for multiple target genes in Arabidopsis in a single generation. Genome Biol 16: 144

Weber E, Engler C, Gruetzner R, Werner S (2011) A modular cloning system for standardized assembly of multigene constructs. PLoS ONE 6: e16765

Werner S, Engler C, Weber E, Gruetzner R, Marillonnet S (2012) Fast track assembly of multigene constructs using Golden Gate cloning and the MoClo system. Bioengineered 3: 38–43

White FF, Taylor BH, Huffman GA, Gordon MP, Nester EW (1985) Molecular and genetic analysis of the transferred DNAregions of the root-inducing plasmid of Agrobacterium rhizogenes. J Bacteriol 164: 33–44

Wolter F, Klemm J, Puchta H (2018) Efficient in planta gene targeting in Arabidopsis using egg-cell specific expression of the Cas9 nuclease of Staphylococcus aureus. Plant J 94: 735–746

Yan L, Wei S, Wu Y, Hu R, Li H, Yang W, Xie Q (2015) High-Efficiency Genome Editing in Arabidopsis Using YAO Promoter-Driven CRISPR/Cas9 System. Mol Plant 8: 1820–1823

Zhang F, LeBlanc C, Irish VF, Jacob Y (2017) Rapid and efficient CRISPR/Cas9 gene editing in Citrus using the YAO promoter. Plant Cell Rep 36: 1883–1887

